# Rational Design of a Protein Kinase A Nuclear-cytosol Translocation Reporter

**DOI:** 10.1101/709766

**Authors:** Allen K. Kim, Helen D. Wu, Takanari Inoue

## Abstract

Protein Kinase A (PKA) exists as a tetrameric holoenzyme which activates with increase of cAMP and plays an important role in many physiological processes including cardiac physiology, neuronal development, and adipocyte function. Although this kinase has been the subject of numerous biosensor designs, a single-fluorophore reporter that performs comparably to Förster resonance energy transfer (FRET) has not yet been reported. Here, we have used basic observations of electrostatic interactions in PKA substrate recognition mechanism and nucleus localization sequence motif to design a phosphorylation switch that shuttles between the cytosol and the nucleus, a strategy that should be generalizable to all basophilic kinases. The resulting reporter yielded comparable kinetics and dynamic range to the PKA FRET reporter, AKAR3EV. We also performed basic characterization and demonstrated its potential use in monitoring multiple signaling molecules inside cells using basic fluorescence microscopy. Due to the single-fluorophore nature of this reporter, we envision that this could find broad applications in studies involving single cell analysis of PKA activity.

## Introduction

As one of the earlier kinases to be characterized, Protein Kinase A (PKA) has been discovered to play an important role in numerous physiological processes including cardiac physiology^1^, neuronal development^2^, and adipocyte function^3^. The kinase, in its inactive form, exists as a holoenzyme composed of two catalytic subunits and two regulatory subunits. Activation of the kinase is generally understood to occur through the increase of cAMP, which leads to the dissociation of the holoenzyme. Kinase activity can further be spatially regulated through the existence of A-kinase anchoring proteins which anchor the holoenzyme to specific subcellular localizations^4^.

This well-characterized nature of PKA has naturally led to it being used as a platform for a number of interesting concepts for kinase reporters. One of the earliest such designs was the utilization of Förster resonance energy transfer (FRET) to monitor the dissociation of the catalytic and regulatory subunits of the protein, a basic design that was used to demonstrate the localness of cAMP concentration and PKA activity^5^. A more generalizable variant of this design was established by Zhang, et al. which involved creating a chimera that consisted of FRET-compatible fluorescent proteins, a phosphopeptide binding domain, and a short kinase substrate sequence^6^. Upon phosphorylation, the binding affinity between the phosphopeptide binding domain and the substrate sequence dramatically increase, leading to a conformational change, which is then measured through changes in FRET. This design has been the basis for a number of generations of PKA FRET reporters in the literature^7–11^.

One of the main advantages of fluorescent kinase reporters is the ability to analyze kinase activity on a single cell level in real time. Despite this advantage, FRET reporters have the inherent downside of requiring two fluorophores, limiting the ability to monitor multiple kinases simultaneously with most experimental setups. Numerous groups have reported various approaches to overcome this limitation in respect to PKA. These approaches have included using multiple FRET pairs^12^, utilizing polarization of light as a readout^13^, or designing proteins that change subcellular localization in response to phosphorylation^14^.

Our approach here utilizes a similar principle that we reported in a previous study where we designed a membrane-bound phosphorylation switch^15^. In summary, arginines in the P-2 and P-3 positions are important residues for substrate recognition^16,17^. By placing the amino acid sequence for the substrate in tandem with minor modifications, we designed an amino acid sequence that loosely conformed to known consensus nuclear localization sequence (NLS) motifs^18^. This peptide responded to PKA activity by shuttling between the nucleus and the cytosol. We furthermore performed basic optimization and demonstrated its use in simultaneously monitoring two different signals within a cell.

## Methods

### Cell Culture and Transfection

HeLa cells were cultured in DMEM with 10% FBS and routinely passaged. Cells were seeded at a density of 2.0×10^4^ cells/cm^2^ in a 6-well plate containing a 25 mm coverslip one day before transfection. They were transfected with multiple plasmids at an amount of 50 ng each using FugeneHD (Promega) according to the manufacturer’s protocol. On the following day, culture media was replaced with serum-free DMEM in the morning. Imaging took place in the afternoon of the same day.

### Image Acquisition

Images were acquired on an inverted microscope (IX81, Olympus) with a heated incubator that maintained the chamber at 37° and 5% CO_2_ (WELS, Tokai-Hit). Images were acquired for either 15 or 25 minutes at 1-minute intervals. Drugs were added at the 5-minute and 15-minute point. PKA activation was achieved through treatment with 50 μM forskolin and 100 μM IBMX, unless otherwise noted. For the experiment demonstrating reversibility, the media was replaced with new media containing 40 μM H89 at the 15-minute point, as residual forskolin and IBMX led to an incomplete inhibition of PKA. Images were acquired with a CMOS camera (C11440, Hamamatsu) in a 60× oil objective (Plan Apo N, Olympus) with the appropriate excitation and emission filters driven by a filter wheel controller. Metamorph was used to control the hardware associated with the microscope which includes the motorized stage (MS-2000, Applied Scientific Instrumentation), filter wheels (Lambda 10-3, Sutter Instruments), and LED light source (pE-300, CoolLED). To measure response with the FRET reporter, cells were excited on the CFP channel, and images were acquired on the YFP channel.

### Image Analysis

All cells were co-transfected with H2B-mCherry (Figs. 1, 2, 3, S1) or H2B-mCerulean3 (Figs. 4, S2, S3, S4) to mark the nucleus. Using the H2B as a mask, ImageJ was used to distinguish cytosolic fluorescent signal from the nuclear signal. The fluorescent signal change was tracked by creating a ratio of cytosol-to-nuclear signal. For FRET experiments, the fluorescent signal was tracked by tracking the ratio from YFP emission to CFP emission. For response profiles, data was normalized by the average of the pre-treatment signals. For all experiments excluding the experiment involving duplex monitoring, analysis was carried out across 3 independent experiments. Ten cells were analyzed from each experiment, resulting in a total of 30 cells analyzed. Student’s t-test for unequal variance was used to test for statistical significance in Figs. 1D and 2D.

**Figure 1:**
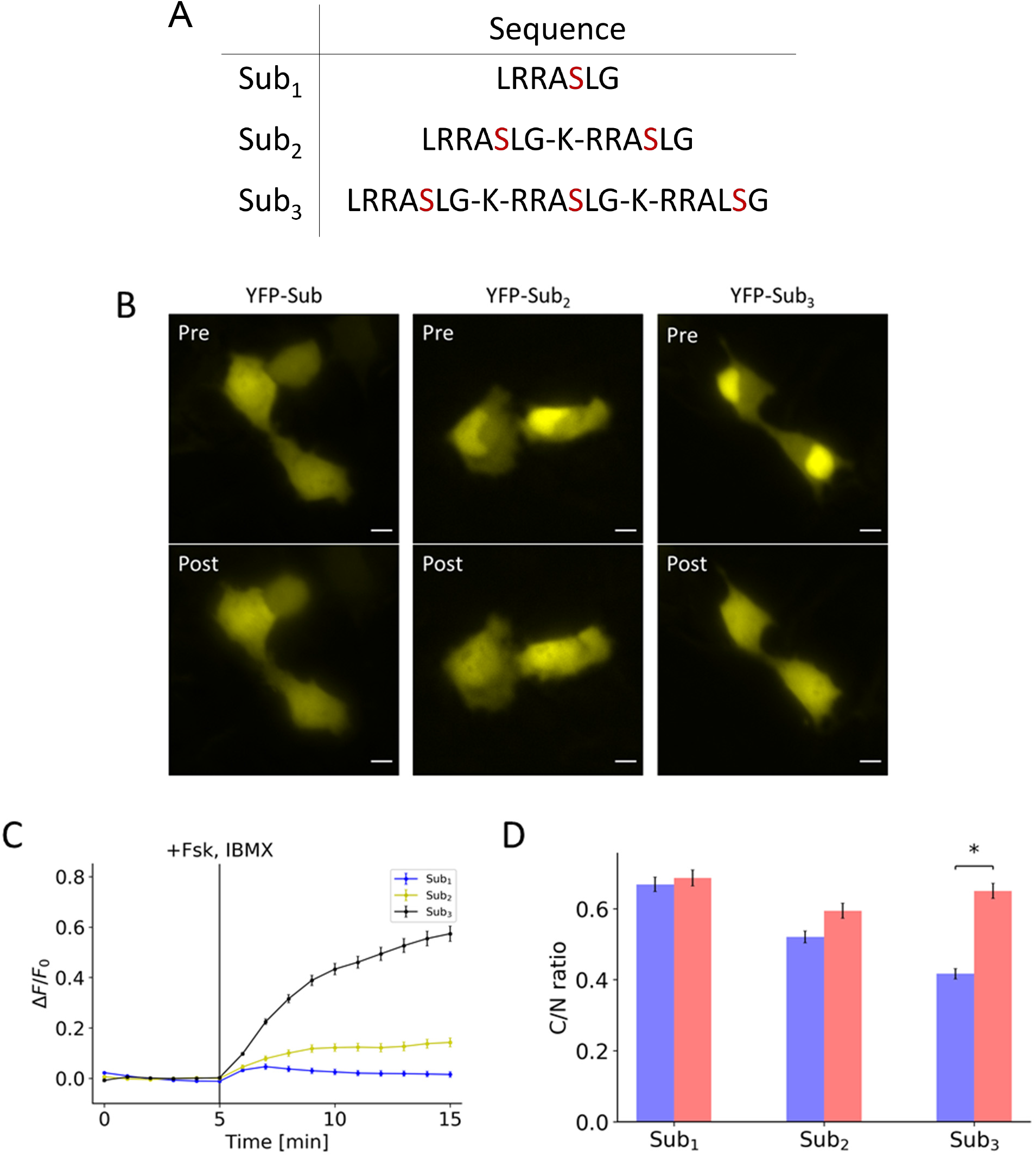
Effects of valency on initial localization and response to PKA activation. **(A)** Table contains the amino acid sequences of the peptides containing the different number of substrates. **(B)** Representative images show the initial distribution (Pre) and the distribution 10 minutes post-treatment (Post) for peptides containing one, two, and three substrates. **(C)** Response profile shows the normalized signal change resulting from the activation of PKA for peptide consisting of one (blue), two (yellow), and three (black) substrates. **(D)** Bar chart represents the pre-treatment localization (blue) and post-treatment localization (red) as measured by taking the ratio of cytosol-to-nuclear fluorescent signal for peptide consisting of one, two, and three substrates. All data points in this figure represent an average signal intensity calculated from 30 cells over 3 independent experiments. Error bar represents standard error of mean. * represents p<0.001. Scale bar represents 10 μm.

**Figure 2:**
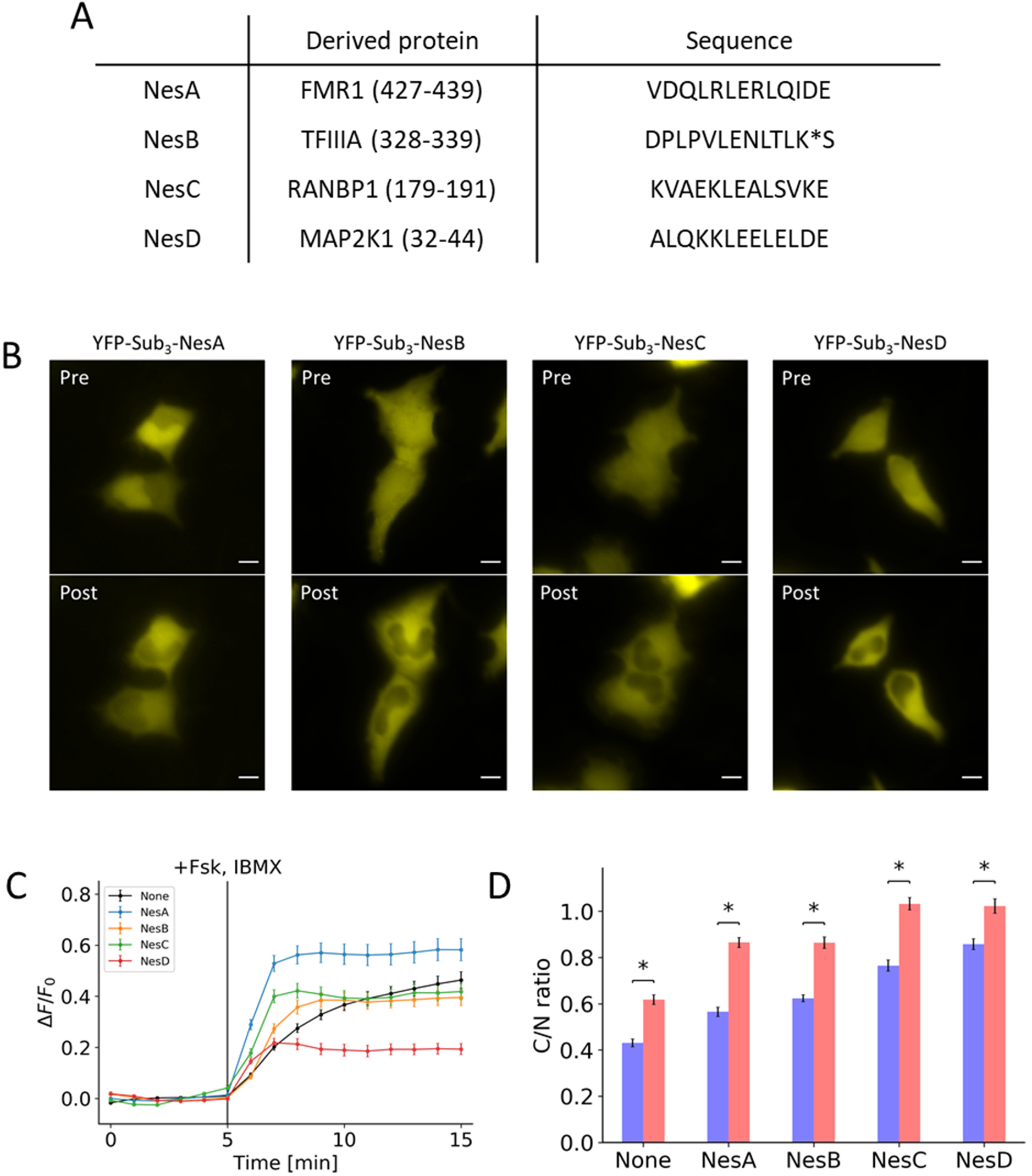
Effects of addition of NES on initial localization and response to PKA activation. (A) Table contains the amino acid sequences of the NES that was fused to the peptide along with the derived gene names. (B) Representative images show the initial distribution (Pre) and the distribution 10 minutes post-treatment (Post) for peptides fused with NesA, NesB, NesC, and NesD. (C) Response profile shows the normalized signal change resulting from the activation of PKA for peptide consisting fused with NesA (blue), NesB (yellow), NesC (green), and NesD (red) along with a peptide without NES (black). (D) Bar chart represents the pre-treatment localization (blue) and post-treatment localization (red) as measured by taking the ratio of cytosol-to-nuclear fluorescent signal for peptides fused with NesA, NesB, NesC, and NesD compared to a peptide without NES. All data points in this figure represent an average signal intensity calculated from 30 cells over 3 independent experiments. Error bar represents standard error of mean. * represents p<0.001. Scale bar represents 10 μm.

**Figure 3:**
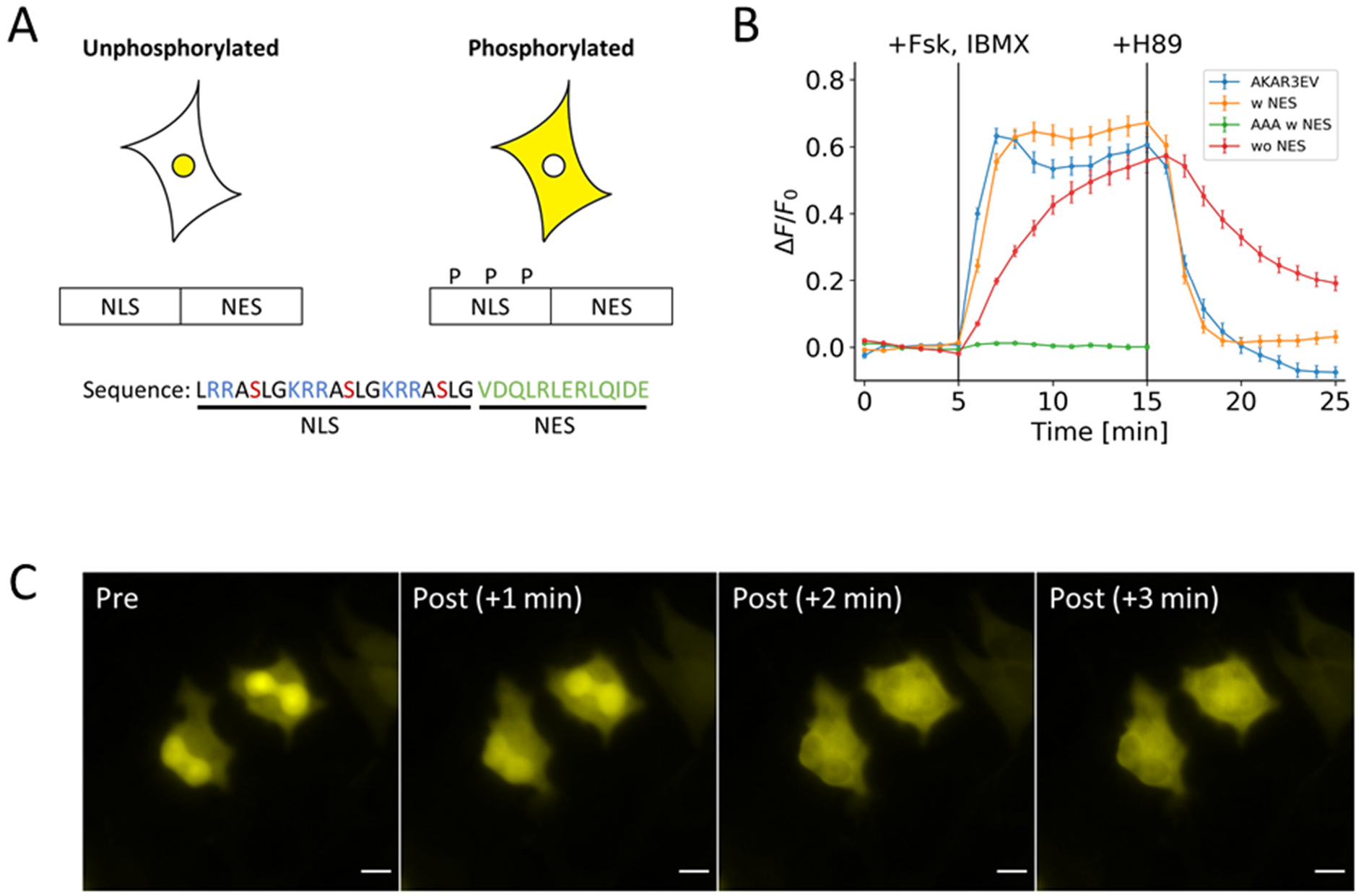
Basic design of the PKA kinase reporter. **(A)** Design consists of a NLS with three phosphorylation sites fused to a NES sequence. Phosphorylation regulates the activity of the NLS. Basic residues are colored blue; phosphoserine is colored red; and NES is colored green. **(B)** Response profile shows the normalized signal change resulting from the activation and inhibition of PKA FRET reporter (blue) and three variants of the nuclear-cytosol translocation reporter. The three variants are: a full version with the NES (yellow), a version lacking the NES (green), and a full version with serine-to-alanine mutations (red). **(C)** Representative images show the changes in distribution of the full version of the reporter that occurs within three minutes of PKA activation. All data points in this figure represent an average signal intensity calculated from 30 cells over 3 independent experiments. Error bar represents standard error of mean. Scale bar represents 10 μm.

**Figure 4:**
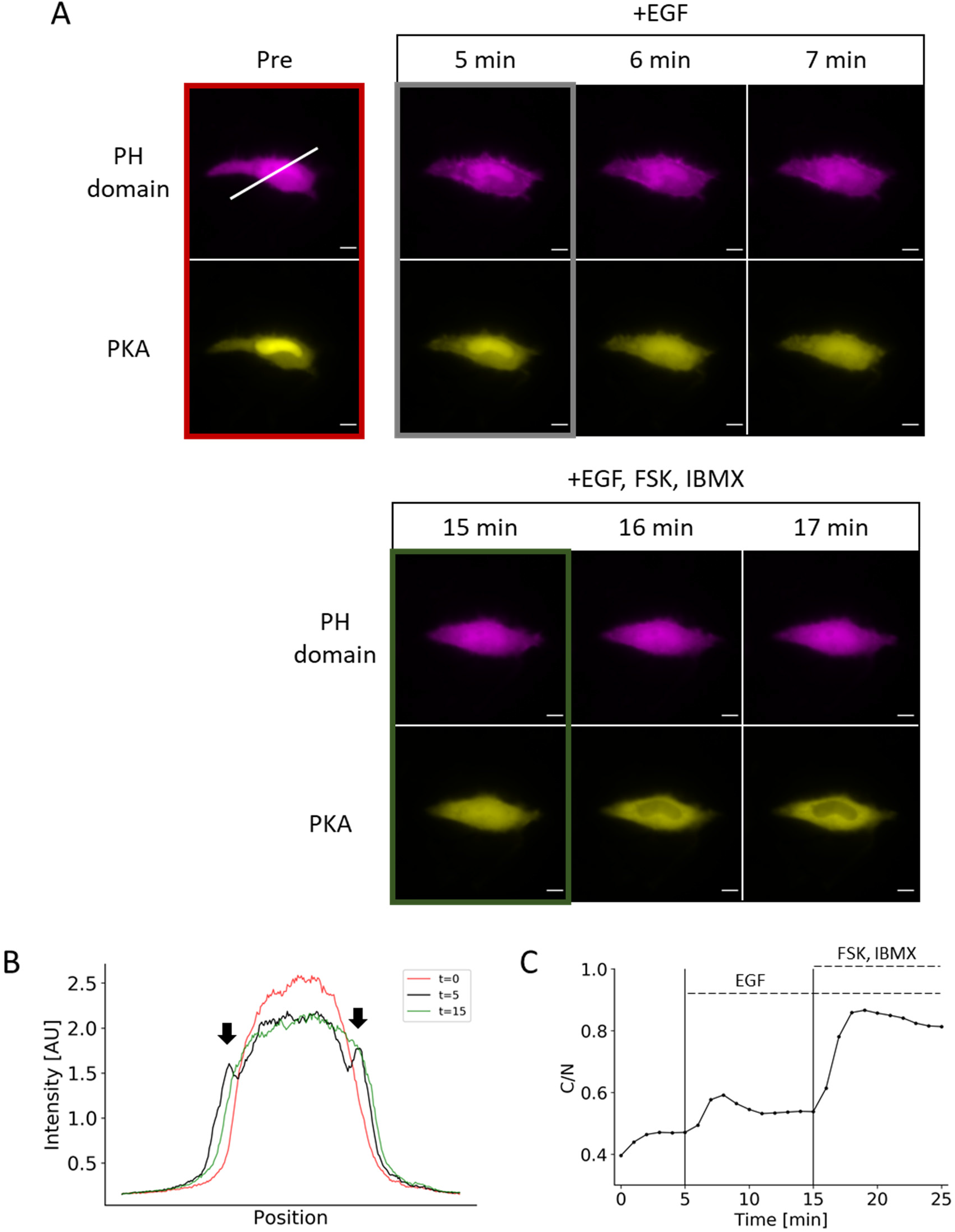
Duplex monitoring of PIP_3_ and PKA in a single cell. (A) Representative images show PIP_3_ accumulation (as indicated by translocation of PH domain) and PKA activity in a single cell after EGF treatment followed by PKA activation. (B) Line-scan of the image in Figure 4A (white line) shows the transient appearance of enriched fluorescent signal at the plasma membrane (arrows). (C) Response profile shows the normalized signal change that results from EGF stimulation followed by a saturation of the reporter response with forskolin and IBMX cocktail. Data points represent measurement from a single cell. Scale bar represents 10 μm.

### DNA Cloning

The cDNA for H2B (gift from Sergi Regot) was amplified by PCR and inserted into NheI and AgeI sites of the mCherry-C1 and mCerulean3-C1 vectors (Clontech). YFP-PH(Akt) is a standard PH domain construct for PIP_3_ labeling. All the reporter variants were inserted into the EYFP-C1 vector (Clontech) at the SacII and BamHI restriction sites using annealed oligonucleotides. The forward sequences of the annealed oligonucleotides are tabulated below:

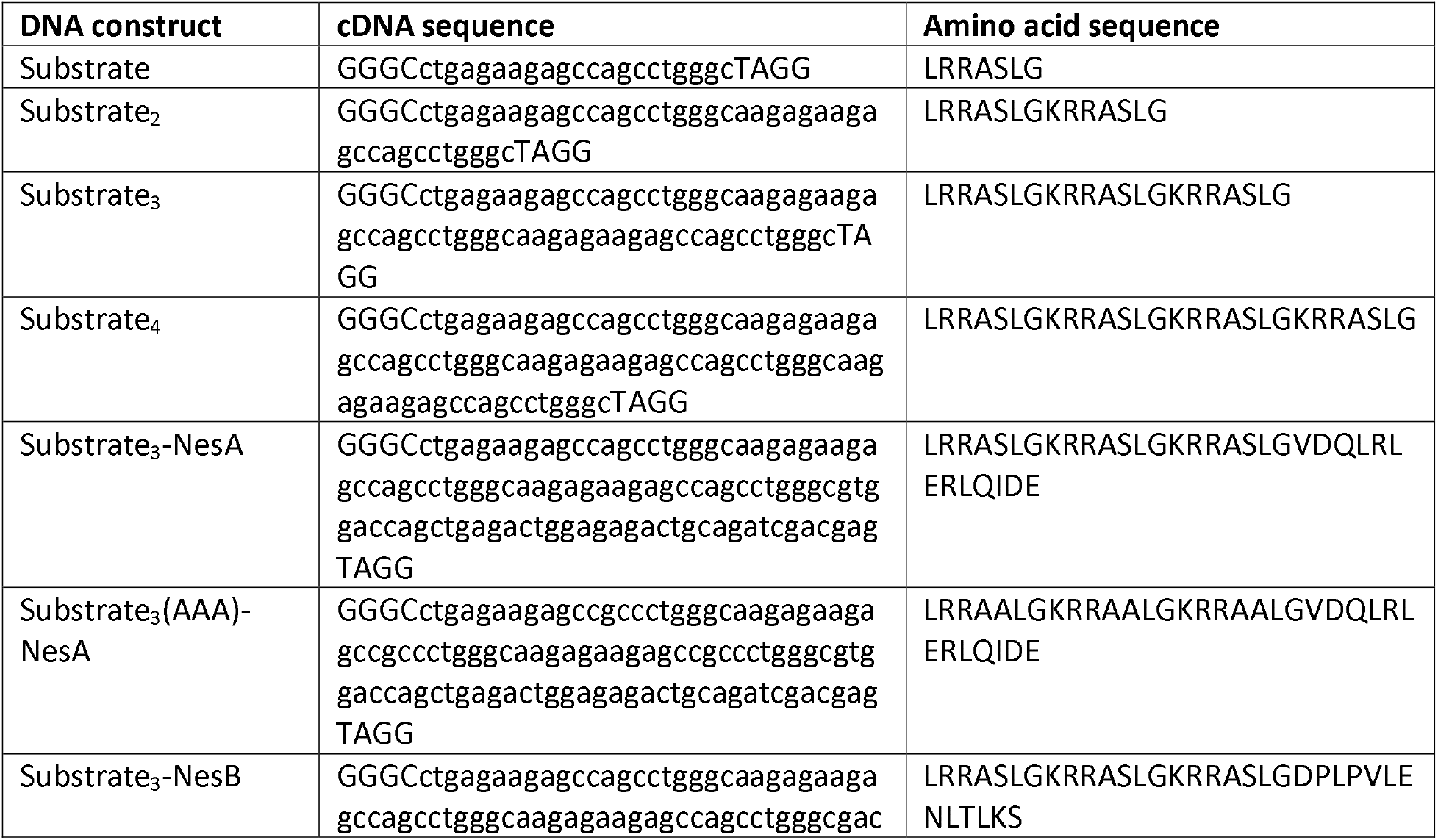
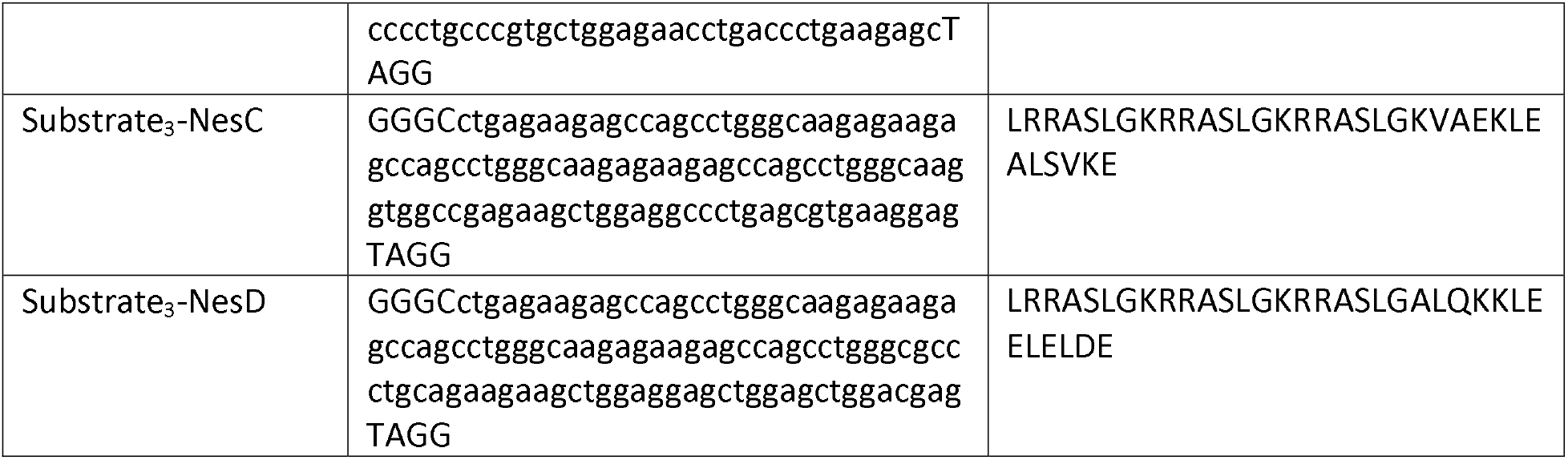

## Results

In an attempt to generate a phosphorylation switch, we hypothesized that the PKA substrate can be arranged in tandem to form a nuclear localization sequence (NLS). Due to the overlapping nature of the NLS with the phosphate acceptor, phosphorylation of this peptide may result in a shift in localization, thus functioning as a switch. To test this idea, we utilized the Kemptide (LRRASLG)^16^, a well-established PKA substrate, as the working unit of the design, which contains the two arginines in the P-2 and P-3 position considered important for substrate recognition. A monovalent substrate alone does not conform to known NLS. However, placing these sequences in tandem leads to a sequence that conforms loosely to NLS motif (Fig. 1A). We transfected cells with EYFP fused to various numbers of substrates. Increasing the number of substrates led to a concomitant increase in nuclear localization (Fig. 1B). When cells were treated with forskolin and IBMX, the fluorescent signal redistributed to become more diffuse at the end of 10 minutes of treatment determined by a normalized response profile (Fig. 1C). The pre-treatment and post-treatment localizations were determined by measuring the ratio of cytosolic-to-nuclear fluorescent signal (Fig. 1D). We also tested a construct containing four substrates. However, because of the strong nuclear localization, there was very little measurable cytosolic signal. For this peptide, we measured the depletion of nuclear signal and observed little difference compared to the construct containing three substrates (Fig. S1). Given these considerations, the trivalent form was considered optimal.

To adjust response kinetics, we next fused a nuclear export sequence (NES) to the trivalent substrate. A previous study established the relative strengths of the various NES’s^19^. Using this as a reference, we fused a 13-amino-acid, leucine-rich fragment from FMR1, TFIIIA, RanBP1, and Map2K1 (Fig. 2A). We observed that three of the four NES’s overpowered the NLS (Fig. 2B). This also translated to a decrease in both response amplitude and kinetics (Figs. 2C, 2D). Amongst the four sequences tested, the NES fragment from FMR1 protein led to a substantial improvement in kinetics and dynamic range in comparison to the peptide lacking a NES.

Based on this characterization, our final design for the PKA phosphorylation switch consisted of NES from FMR1 and the three phosphate acceptors for PKA that functions as an NLS when non-phosphorylated (Fig. 3A). When this optimized switch was fused to EYFP and transfected in cells, this peptide responded to PKA activation by 50 μM forskolin and 100 μM IBMX by redistributing from the nucleus to the cytosol (orange line, Fig. 3B). When the signal change was quantified by taking the ratio of cytosolic fluorescence signal to nuclear signal, the kinetics and dynamic range was comparable to the FRET reporter, AKAR3EV^9^ (blue line, Fig. 3B). The response to 40 μM H89, which resulted in the inhibition of PKA, also revealed similar kinetics and response to its FRET counterpart. A serine-to-alanine mutation of all the phosphate acceptors led to no response (green line, Fig. 3B), while removing the NES led to a much slower response to activation (red line, Fig. 3B). Representative images show the change in localization of the EFYP within the 3 minutes following PKA activation (Fig. 3C). To further validate that PKA is the kinase that phosphorylates the PKA switch, we used other PKA inhibitors, namely Rp-cAMPS and PKA peptide inhibitor (PKI). When cells were treated with 500 μM Rp-cAMPS following IBMX/Fsk stimulation, the PKA switch altered its subcellular localization (Fig. S2A). We observed almost complete suppression of PKA switch response when the IBMX/Fsk cocktail was added to cells expressing PKI fused to mCherry, while this suppression was not present in cells expressing mCherry alone (Figs. S2B, S2C). These data clearly support that our switch peptide is controlled by PKA activity.

One of the advantages of the PKA phosphorylation switch is a potential for multiplex imaging. To demonstrate this, we stimulated cells with two different ligands, EGF and isoprenaline. First, by co-transfecting the PH domain from Akt (mCherry) and the PKA phosphorylation switch (EYFP), we simultaneously monitored the synthesis of PIP_3_, a lipid secondary messenger, and the activity of PKA. EGF activates multiple signal pathways in parallel, leading mainly to PI3K-mediated PIP_3_ production and MAPK activation, and to PKA activation to a lesser extent in some cell types^20^. Upon stimulation of HeLa cells with 50 ng/mL EGF, we observed a transient increase in PIP_3_, indicated by the translocation of the PH domain from the cytosol to the plasma membrane (Fig. 4A). This was indicated by the accumulation of PH domain at the plasma membrane upon activation with EGF but not upon PKA activation (Fig. 4B). PKA activity as measured by the reporter led to a sustained level of increase upon activation of EGF and saturated with treatment of the forskolin and IBMX cocktail (Fig. 4C). We also reversed the order of treatment and observed PIP_3_ enrichment only when the cells were treated with EGF (Fig. S3). Second, we added 10 μM isoprenaline to cells that have been co-transfected with the PKA phosphorylation switch and R-GECO1, intending to measure PKA activity and calcium ions (Ca^2+^), respectively. As a result, we obtained a distinct temporal response of these two signaling molecules; while Ca^2+^ indicated a sharp and transient increase (Fig. S4A), PKA activity exhibited a more sustained increase (Fig. S4B). Together, the duplex monitoring of two signal activities was achieved in live cells using the PKA phosphorylation reporter (Fig. 4 and Fig. S4C).

## Discussion

Our PKA phosphorylation switch is a *de novo* amino acid sequence based on simple observations of PKA consensus sequence and NLS motif. Using this information, we designed a switch that reproduced the kinetics measured with a well-established FRET reporter. In contrast to FRET reporters, this type of single-fluorophore reporter frees up a fluorescent color, allowing for an additional channel of analysis. We then demonstrated this capability monitoring two different signaling molecules by overexpressing cells with the PH domain from Akt and the PKA reporter.

A switchable NLS alone did not reproduce the kinetics of PKA activation observed with a FRET reporter. The distribution of a protein observed in a cell is determined by the net flux of the nuclear import and nuclear export transport processes. With peptides containing only NLS’s, the nuclear import is driven by facilitated diffusion while nuclear export is driven by passive diffusion. As passive diffusion is slower than facilitated diffusion, the time required to reach steady state is longer, reflected by the decrease in response kinetics. The addition of a NES mitigated this issue. An ideal reporter, thus, should be rate-limited by the kinetics of the underlying phosphorylation, not by auxiliary processes.

Another important property of reporters is kinase specificity. In our study, we have used a well-characterized substrate sequence of PKA that has been used in previous FRET reporters^7–11^. A phosphorylation prediction algorithm, NetPhos3.1^21^, strongly suggests PKA as the target kinase for this reporter. Furthermore, based on the current literature of AGC kinases, PKA is the only AGC kinase that requires two arginines at P-2 and P-3 position^16,17,22,23^. These reports do not rigorously rule out non-specific phosphorylation, as substrates are not necessarily required to conform to consensus sequences. However, given current literature and our data, PKA activity appears to be the driving force behind the shuttling.

We used the interaction of positively charged residues in the substrate sequence with the catalytic subunit of PKA to design the reporter. However, this is not necessarily the only method of achieving specificity. Regot *et al.* developed a similar reporter which shuttles between the cytosol and the nucleus using a different strategy^14^. The phosphate acceptors in this reporter are placed in a much weaker, proline-rich substrate recognition sequence derived from c-Jun. The affinity towards the substrate is increased through docking sequences, which elevates the effective concentration of the substrate to the kinase^24,25^. We were unable to observe a measurable response with this PKA reporter. However, we speculate that this reporter design relies on a much weaker interaction with the kinase than our design, requiring a more careful optimization of experimental conditions.

In summary, we have designed a reporter based on a rearrangement of a PKA substrate to create a NLS. In principle, this is generalizable to all basophilic kinases, as this mechanistically relies on disruption of electrostatic interactions through phosphorylation. Due to the single-fluorophore nature of this reporter and its comparable performance to a FRET reporter, we envision that this could find broad applications in studies involving single cell analysis of PKA activity.

## Acknowledgments

We would like to thank Dr. Kazuhiro Aoki for providing the AKAR3EV construct. This study was supported by National Institute of Health R01 GM123130 and DARPA HR0011-16-C-0139 (to T.I.).

## Author Contributions

T.I. and A.K.K. conceived and designed this study. A.K.K. and H.D.W. conducted experiments and analyses. A.K.K. wrote the manuscript with contributions from T.I.

