## Supplementary Information for "Rational Design of a Protein Kinase A Nuclear-cytosol Translocation Reporter"

Content: Figure S1-S4

Figure S1

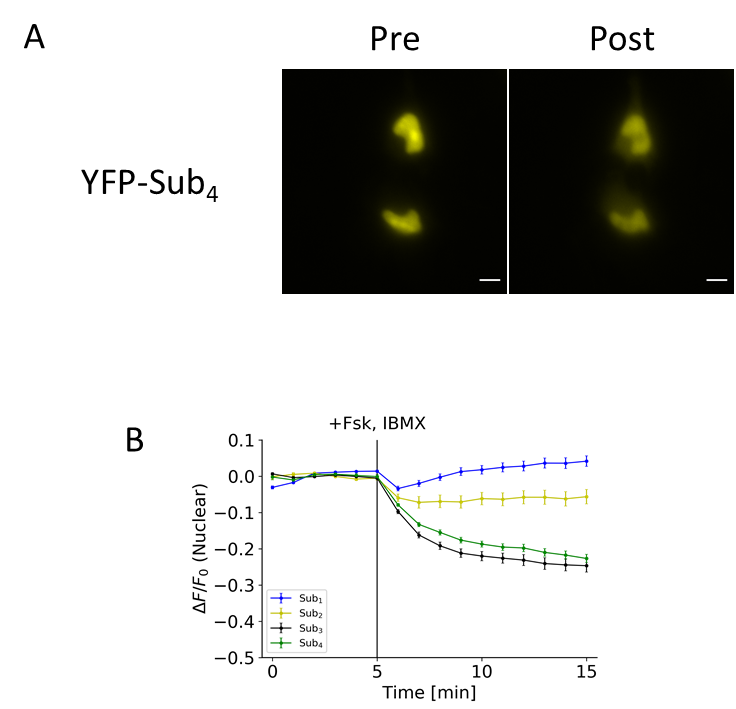

**Figure S1: Response of tetravalent substrate to PKA activation.**

(A) Representative images show the localization of YFP-Sub_4_ pre-treatment (Pre) and 10 minutes post-treatment (Post). Sub_4_ peptide sequence is following: LRRASLG-K-RRASLG-K-RRASLG-K-RRASLG.
(B) Response profile shows the normalized change in nuclear fluorescent signal after PKA activation with the peptides containing the different number of substrates.

All data points in this figure represent the average signal intensity calculated from 30 cells over 3 independent experiments. Error bar represents standard error of mean. Scale bar represents 10 µm.

Figure S2

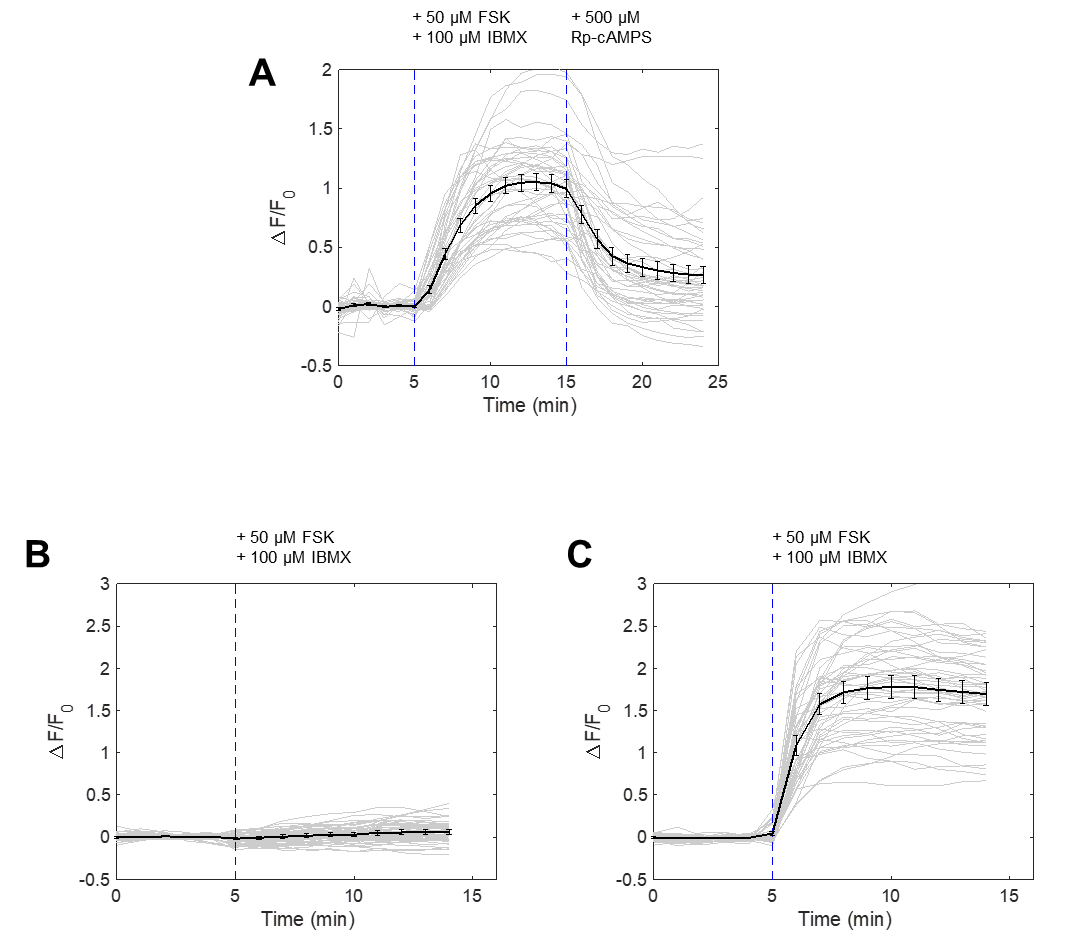

**Figure S2: Specificity of PKA kinase reporter to PKA activation by pharmacological and genetic inhibition.**

(A) Nuclear-cytosol translocation of PKA kinase reporter by FSK/IBMX treatment and reversal with PKA inhibitor Rp-cAMPS (500 μM, Santa-Cruz, Sc-24010). (N = 3 experiments, 45 cells)
(B) Nuclear-cytosol translocation of PKA kinase reporter in cells co-expressing mCherry-PKI. (N = 3 experiments, 54 cells)
(C) Nuclear-cytosol translocation of PKA kinase reporter in cells co-expressing mCherry. (N = 3 experiments, 51 cells)

### Figure S3

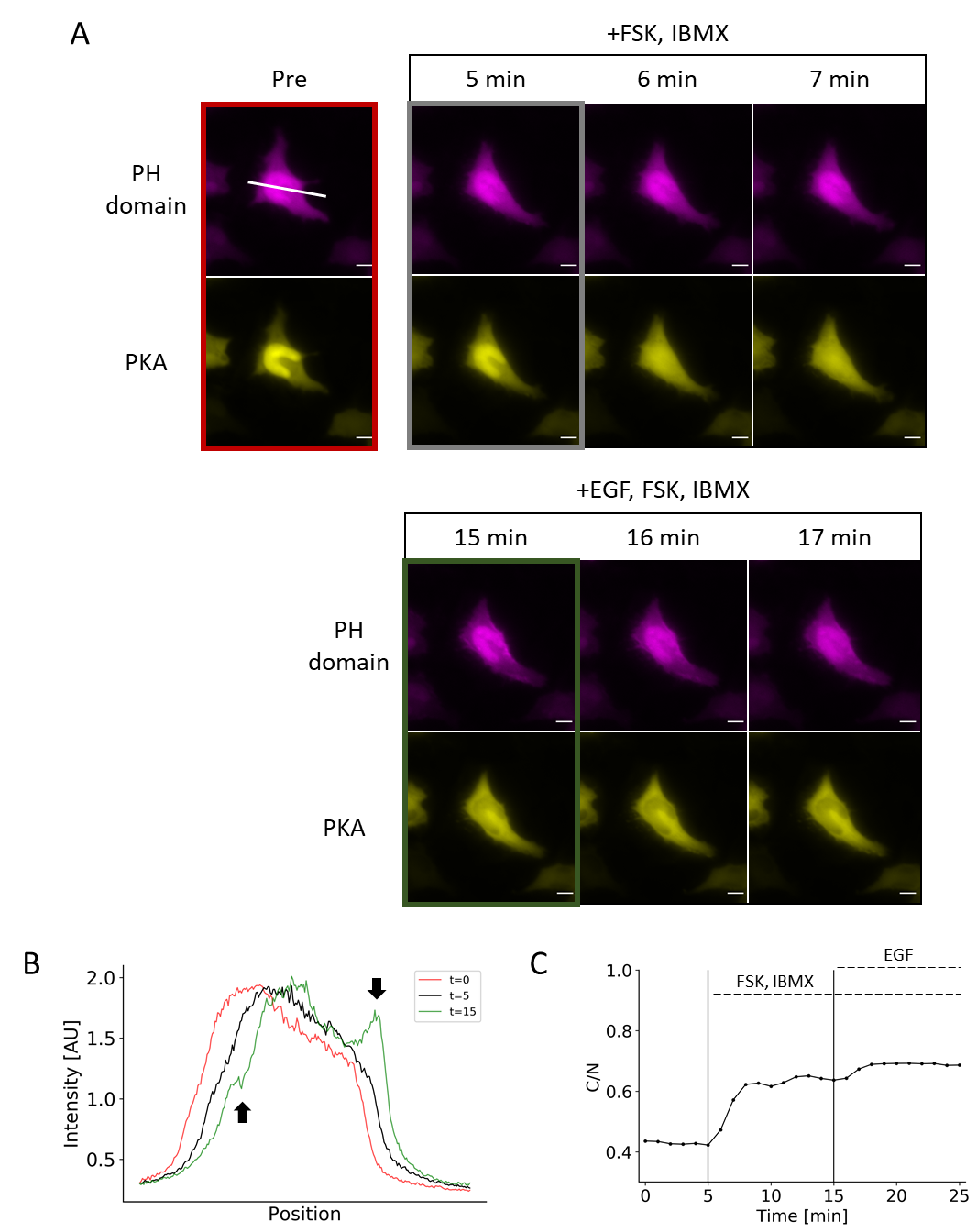

**Figure S3:** **Duplex monitoring of PIP_3_ and PKA in a single cell with PKA activation followed by EGF treatment.**

(A) Representative images show PKA activity and PIP_3_ accumulation (as indicated by translocation of PH domain) in a single cell after PKA activation followed by EGF treatment.
(B) Line-scan of the image in Figure 4A (white line) shows the transient appearance of enriched fluorescent signal at the plasma membrane with EGF treatment but not with PKA activation (arrows).
(C) Response profile shows the normalized signal change that results from PKA activation followed by EGF stimulation.

Data points represent measurement from a single cell. Scale bar represents 10 µm.

Figure S4

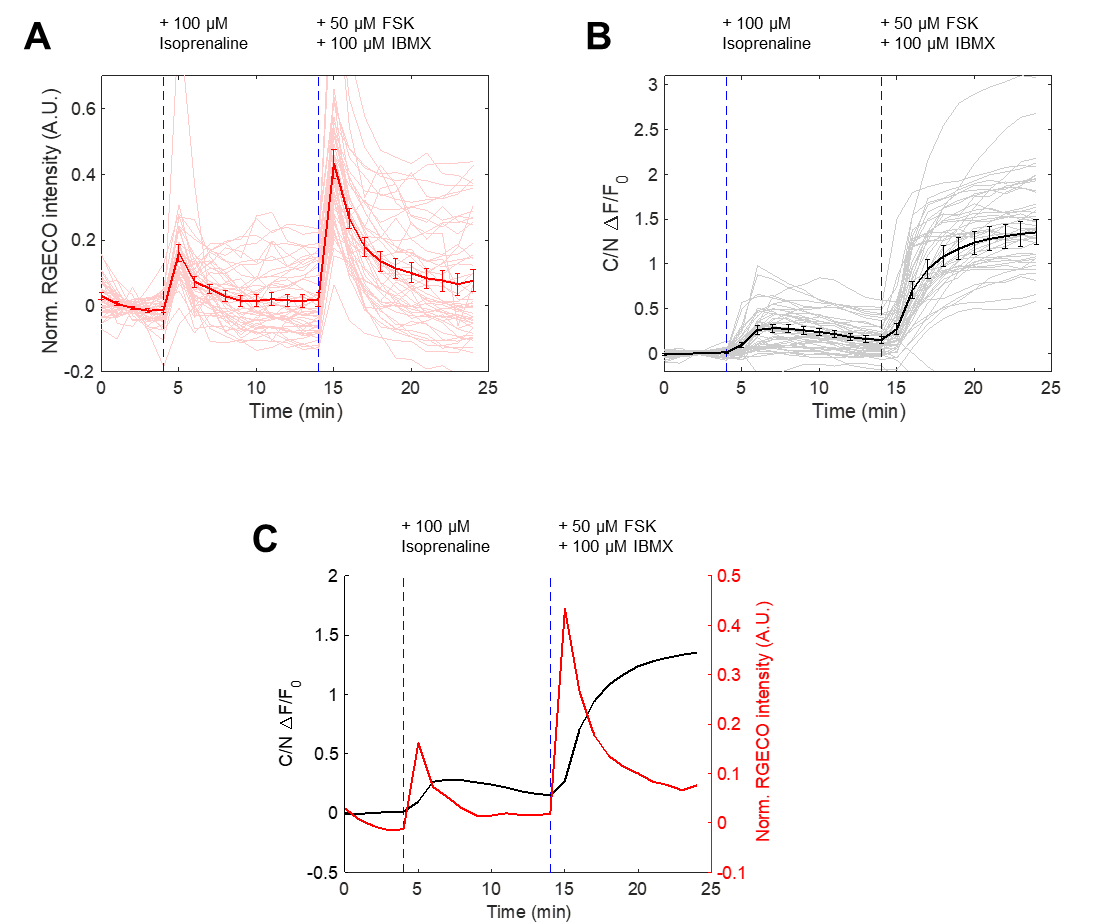

**Figure S4: Duplex monitoring of Ca2+ and PKA in cells followed by isoprenaline and FSK/IBMX treatment.**

(A) Ca^2+^ response as reported by R-GECO to isoprenaline (100 μM, Sigma-Aldrich, I5627-5G) and FSK/IBMX treatment in cells also expressing PKA kinase reporter and nucleus marker. (N = 3 experiments, 48 cells)
(B) Nuclear-cytosol translocation response of PKA kinase reporter in same cells in (A).
(C) Duplex monitoring of Ca^2+^ and PKA activity from (A) and (B).
